# Does genetic rescue disrupt local adaptation? An experimental test using thermally adapted *Tribolium castaneum* lines

**DOI:** 10.1101/2025.08.07.669094

**Authors:** George West, Michael Pointer, Will Nash, Rebecca Lewis, David S Richardson

## Abstract

Anthropogenic drivers are restricting many species to small, genetically isolated populations. These are prone to inbreeding depression and are at an increased risk of extinction. Genetic rescue, the controlled introduction of genetic variation from another population, can alleviate inbreeding effects. A major conservation concern, restricting the use of this technique, is that such augmented gene flow may disrupt local adaptation crucial to a population’s persistence. Using populations of the red flour beetle (*Tribolium castaneum*) experimentally adapted to reproduce at higher temperatures, we assess whether genetic rescue attempts disrupt thermal adaptation. Rescuers, drawn from populations adapted to either 30°C or 38°C, were introduced into populations adapted to 38°C, which had been inbred for two generations. We recorded population productivity for three generations post-rescue, in the adapted 38°C environment. Rescuers with and without local adaptation significantly increased the productivity of recipient inbred populations but, importantly, those sharing local adaptation to reproduction at 38°C provided greater increases in productivity. For the first time, we show that co-adaptation between rescuing individuals and rescuee population maybe an essential aspect of achieving desired conservation outcomes.

## Introduction

Climate change and habitat destruction are fragmenting species into increasingly small and isolated populations where gene flow is disrupted. This results in inbreeding, inbreeding depression and, consequently, increased risk of extinction (Haddad et al., 2015). Inbred individuals are more likely to be homozygous for any recessive, deleterious alleles present in the population, exposing their harmful effects (Crnokrak and Roff, 1998; Charlesworth and Willis, 2009). This conversion of hidden genetic load, masked by dominance, into expressed load contributes to reduction in fitness at both the individual- and population-level. This increase in homozygosity may also lead such individuals to lose the benefits of heterozygote advantage (Hedrick and Garcia-Dorado, 2016). Factors such as these interact with environmental drivers to push populations towards extinction (Soule and Gilpin, 1986; Blomqvist et al., 2010; Palomares et al., 2012).

Genetic rescue refers to the increase in fitness observed in an inbred population when novel genetic variation is introduced by a conspecific from another population, a ‘rescuer’ (Ingvarsson, 2001; Hedrick et al., 2011). This process increases genome-wide heterozygosity within the target population, reducing expressed genetic load and improving fitness. Genetic rescue has been successfully implemented in endangered populations ( Clarke et al., 2024; Frankham, 2016, 2015; White et al., 2023), including the Florida panther (*Puma concolor couguar*) (Pimm et al., 2006; Onorato et al., 2024). Experimental systems have also extended our understanding of genetic rescue (Hufbauer et al., 2015; Fitzpatrick et al., 2019; Zajitschek et al., 2009; Bijlsma et al., 2010). Despite this body of evidence supporting its efficacy, genetic rescue remains controversial among conservation managers (Frankham et al., 2011; Edmands, 2007).

If populations in need of genetic rescue exhibit local adaptation, the input of novel genetic variation from other populations could disrupt beneficial gene complexes. Rescuers may introduce non-locally adapted alleles, the expression of which could disrupt adaptation, and thus exacerbate reduced fitness in vulnerable recipient populations (Kawecki and Ebert, 2004; Lenormand, 2002). This effect, termed outbreeding depression, has been suggested to be a key risk of implementing genetic rescue as a conservation measure (Bell et al., 2019; Tallmon et al., 2004). However, there are few examples of outbreeding depression (Hedrick et al., 2019; Turček and Hickey, 1951; Loope et al., 2024), and it has been argued that, if genetic rescue guidelines are followed, the risk of such a detrimental impact is overstated (Frankham et al., 2011; Ralls et al., 2020; Powell, 2023), relative to the potential benefits of rescue.

Selecting the source population for genetic rescue attempts is key to avoiding outbreeding depression. Populations from across a species range may be locally adapted to different conditions, leading to the introduction of maladaptive traits reducing population fitness (Bachmann et al., 2020). Increasingly reintroductions from captivity are being considered to reinforce wild populations including in the context of genetic rescue (Al Hikmani et al., 2024). Captivity, however, could promote maladaptation as natural selection is weakened if not absent (Frankham, 2008), utilising individuals from captivity for rescue could reduce the fitness of wild populations by introducing maladapted genotypes. Testing the effects of maladaptation on genetic rescue, from wild or captive populations, is vital to improve our ability to select the best source population for rescuers.

Climate change poses a significant challenge endangered species (Hoegh-Guldberg et al., 2018), with genetically depauperate populations struggling to adapt (Bellard et al., 2012). The introduction of genetic variation into isolated populations via genetic rescue could expedite adaptation to rapidly evolving climate by supplementing standing genetic variation (Bell and Gonzalez, 2009; Bell et al., 2019). Seeding rescue attempts from populations with specific local adaptations, for example to high temperatures (Macadam et al., 2025), could facilitate the introgression of beneficial adaptations into endangered populations (Kelly and Phillips, 2019; Rudin-Bitterli et al., 2021). Genetic rescue of inbred endangered populations that exhibit specific local adaptations is also a key consideration to protect such unique combinations of alleles towards future species level resiliency.

*Tribolium castaneum,* a Tenebrionid beetle, is a model system (Pointer et al., 2021) for population genetics, genetic rescue, and thermal tolerance (Hufbauer et al., 2015; Sales et al., 2021; Sokal and Sonleitner, 1968). Here, we use *T. castaneum* populations experimentally selected for over 150 generations to reproduce at 38°C, compared to the ancestral population optimum of 30°C (Vasudeva et al., 2019; Skourti et al., 2022). When reared in 30°C conditions, these thermally adapted populations are less fit than replicate populations maintained at 30°C, which produce more offspring when kept as adults at both 30°C and 38°C (Lewis, 2020). However, when reared at 38°C, thermally adapted beetles produce more eggs than non-adapted beetles, and those eggs have greater hatching success (Lewis, 2020). In order to test the impact of genetic rescue from rescuing individuals with and without local adaptations, we used replicated subpopulations of these adapted lines to generate inbred experimental populations. We assessed the impact of genetic rescue from individuals with different adaptive backgrounds on thermal tolerance adaptation. We show that genetic rescue by an adapted individual was the most effective treatment at increasing fitness in inbred populations.

## Methods

### Husbandry

*T. castaneum* populations were maintained on standard fodder (90% white organic flour, 10% brewer’s yeast and a layer of oats for traction) in a controlled environment of 30°C (unless otherwise stated) and 60% humidity with a 12:12 light-dark cycle. Populations were maintained following a standard cycle of virgin adults having seven days of mating and oviposition followed by the removal of adult beetle, using 2 mm and 850 µm sieves, so that only eggs remain in the fodder. Each generation is initiated with a number of adults (line dependent, see below) that are given 7 days to mate and lay eggs before being removed. The eggs are left for 35 days to develop into mature adults.

### Tribolium castaneum lines

#### Krakow super strain (KSS)

A combination of fourteen laboratory strains bred to maximise genetic diversity in one population maintained at a census size of 600 individuals (Laskowski et al., 2015). This line is highly productive at 30°C but has reduced fitness at 38°C. This was used as the non-adapted rescuer population.

#### Thermal lines

Ten independent lines (census size = 100 adults) founded from KSS, and experimentally evolved for ∼150 generations at an environmental temperature of 38°C (Dickinson, 2018), thus imposing selection for development and reproduction at this temperature, considerably above the thermal optimum for *T. castaneum* (Howe, 1962). All other conditions were as described above aside from a shorter development period of 27 days, reflecting accelerated development at 38°C. These were used as the thermally adapted rescuer populations.

#### Inbred lines

Ten inbred populations were created, one from each of the ten thermal lines described above. Adult beetles from each thermal line were housed individually for two weeks to ensure any fertilized eggs were laid. Single-pair matings were formed by housing together a previously isolated male and a female for 7 days of mating and oviposition, resulting in a single pair bottleneck for each thermal line. Full sibling offspring resulting from this pairing were again paired for a second bottleneck. The following generation was initiated with 10 male and 10 female full sibling offspring of full sibling pairs. From the offspring of these groups, six inbred experimental populations (10 males and 10 females) were created from each of the 10 inbred thermal lines, to act as recipient populations for genetic rescue. One inbred population only produced four experimental populations, resulting in a total of 58 experimental populations split over two temporal blocks of 30 and 28. The two blocks were maintained one day apart for ease of handling but otherwise received identical treatment. Each population received a random ID number to blind the experiment and avoid bias. Experimental populations were initiated every generation using 10 males and 10 females sourced from the offspring of the previous generation to reduce density dependent effects (Duval et al., 1939; King and Dawson, 1972; Janus, 1989). All experimental inbred recipient populations were kept at 38°C in A.B. Newlife 75 Mk4 forced air egg incubators (A.B. Incubators, Suffolk, UK); all other conditions were kept as described above.

### Genetic rescue protocol

Populations were kept in 125 ml tubs containing 70 ml of standard fodder. After seven days, adults were discarded and eggs were left to develop for ∼21 days when 10 male and 10 female pupae were randomly selected to establish the next generation. Remaining individuals were maintained for 10 days following this and were then frozen and manually counted. Pupae taken at day ∼21 were housed in plastic dishes containing 10ml standard fodder in single-sex groups until they matured into adults after 10±2 days, and the next generational cycle began with unmated adults, avoiding overlapping generations.

After a rest generation a single male from each inbred recipient population was removed and replaced with a single male rescuer to avoid demographic rescue effects (increased population fitness due to increased population size) (Bell et al., 2019; Ingvarsson, 2001). Three treatments were created: 1) control - 19 populations received no rescue (the male was not removed); 2) locally adapted rescue - 19 populations received a 38°C-adapted rescuer (a male from a different thermally adapted population); 3) non-locally adapted rescue - 20 populations received a non-thermally adapted rescuer (a KSS male, see above) (figure 1).

**Figure 1:**
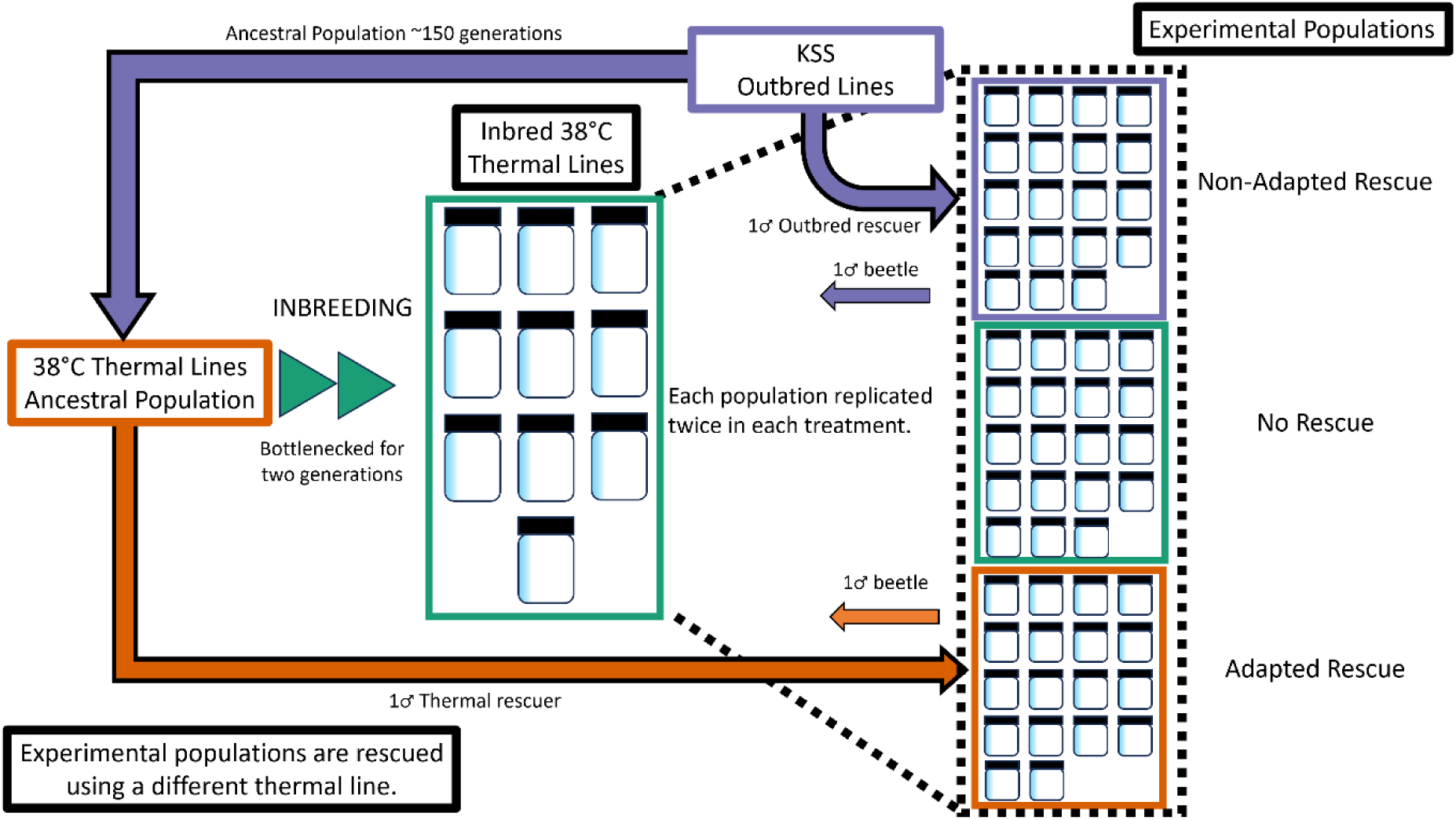
Experimental design for the attempted genetic rescue of inbred thermally adapted *T. castaneum* populations by a single thermally adapted or non-thermally adapted rescuer. Inbred, thermally adapted lines were created by inbreeding lines thermally adapted to 38°C over ∼150 generations (Dickinson, 2018) with two generations of full sibling matings, before being kept for three generation at n = 20 (10 females and 10 males) during the experiment. The 10 inbred thermal lines were replicated to be represented in each experimental treatment twice. The final sample size was 56 experimental populations (see main text).

Population fitness was measured using productivity: the number of mature adult offspring the population produced each generation. Populations were maintained for three non-overlapping generations following rescue. Two replicates were lost after two generations due to human error, resulting in a third generation with 19 control, 18 thermally adapted rescue, and 19 non-thermally adapted rescue populations. Data for populations lost in generation three was included in the analysis of generations one and two.

### Statistical analysis

R v.4.4.1 (R Core Team, 2024) was used with R Studio version 2024.04.2+764 (Posit team, 2024). Data management and exploration were performed with tidyverse (Wickham et al., 2019), stats (R Core Team, 2024), Rmisc (Hope, 2022) and googlesheets4 (Bryan, 2023). ggplot2 (Wickham, 2016) was used to visualise results. Data distribution was checked using the shapiro.test function (R Core Team, 2024). The glmmTMB package (Brooks et al., 2017) was used to fit generalised linear mixed models (GLMMs). DHARMa (Hartig, 2022) was used to check model fit and the check_collinearity function from the performance package (Lüdecke et al., 2021) to test variance inflation factor scores (VIF). No overdispersion or collinearity (VIF<3 for all variables) was found. R^2^ was determined using the r.squaredGLMM function in MuMIn (Bartoń, 2024).

Counts of population productivity over all generations was analysed using a GLMM with a negative binomial errors and a log link function. Fixed explanatory variables were rescue treatment and generation as well as the interaction between these variables. To account for populations variance and relatedness between replicate populations a random factor was added nesting individual ID within the stock thermal line the population descended from. The control treatment was set as the baseline factor for comparison. The baseline was changed to non-adapted to compare between the two rescue treatments post-hoc.

GLMMs, constructed as described above, but excluding the generation variable, were then run post-hoc on each generation individually to test if there were significant differences between the treatments in each generation. The baseline was changed to non-adapted rescuers, to compare between the two rescue treatments post-hoc.

## Results

The interaction between treatment and generation was significant for the thermally-adapted rescue (β = 0.093 ± 0.039, z = 3.38, P = 0.017) but not for the standard rescue (β = 0.055 ± 0.039, z = 1.41, P = 0.16) when each was compared with the no-rescue control (Table 1, Fig. 2). A post-hoc contrast between the two rescue treatments showed no difference in the gradient of productivity change over generation (β = 0.038 ± 0.037, z = 1.01, P = 0.31, 95 % CI = –0.035 – 0.111).

**Figure 2:**
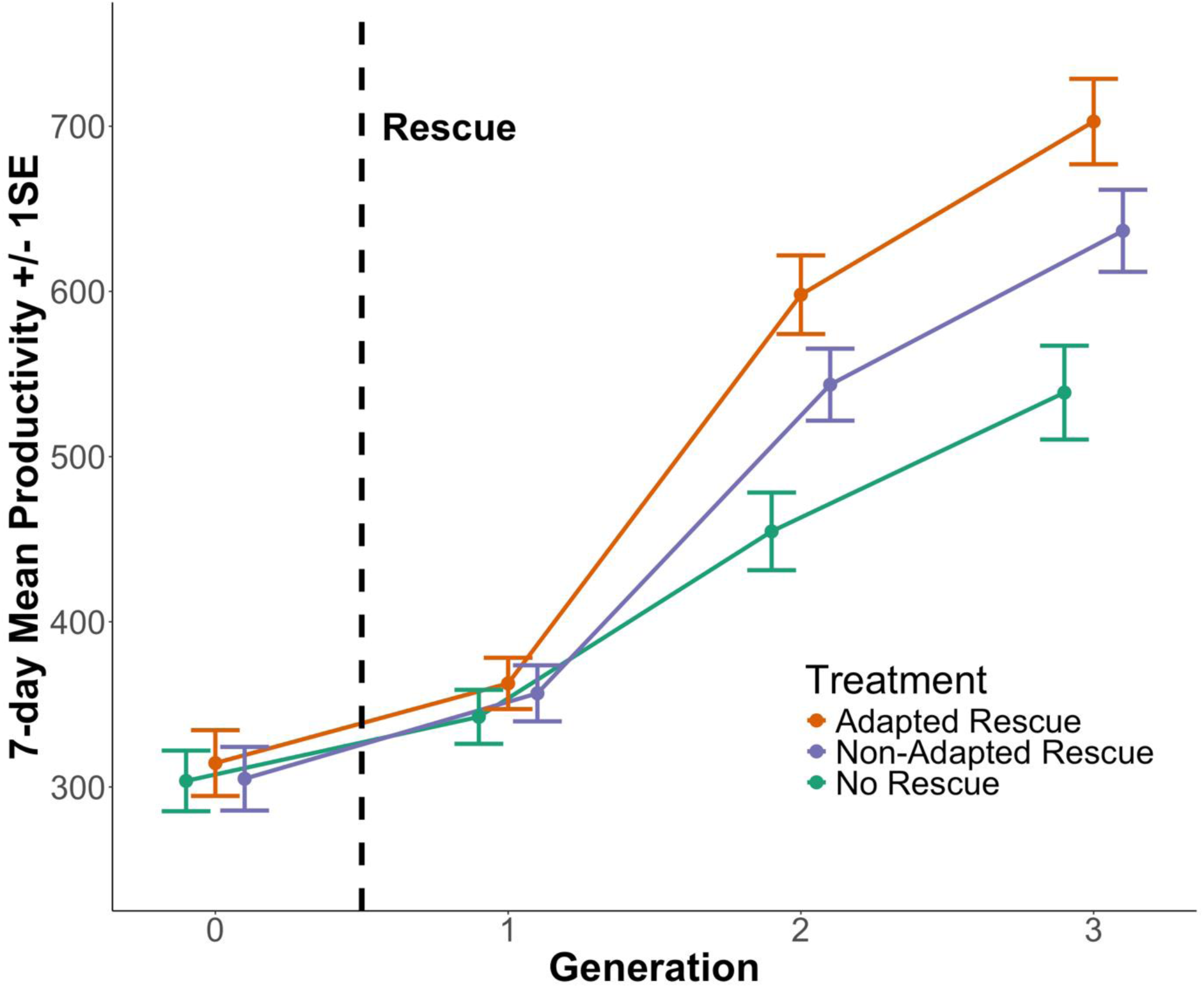
The effect of introducing a 1) thermally adapted (orange) or 2) non-adapted (blue) rescuer, compared to control populations (green) on the mean productivity of inbred thermally adapted populations of *T. castaneum* (Population size = 20, Number of experimental populations = 58 (56 in generation 3)) over three generations after a single introduction event (dotted vertical line) while maintained at 38°C. Generation 0 was not included in statistical analysis. Plot is jittered to aid in visualisation, Error bars represent +/- 1 standard error. Generation 0 not included in analysis.

**Table 1:** Summary of a GLMM fitted to model the productivity of small, inbred, thermally adapted *T. castaneum* populations (Population size = 20, Number of populations = 58) after receiving a rescue by a thermally adapted, or non-adapted, male rescuer or no rescue over three generations. Predictors in bold are significant (*P <* 0.05). Marginal R^2^ = 0.637, Conditional R^2^ = 0.740.

| Predictor | Estimate | SE | z | P |
| --- | --- | --- | --- | --- |
| Intercept | 5.626 | 0.0756 | 47.43 | <2e-16 |
| Treatment (No Rescue) |  |  |  |  |
| Thermally Adapted Rescue | 0.024 | 0.096 | 0.25 | 0.803 |
| Non-adapted Rescue | 0.028 | 0.096 | 0.29 | 0.771 |
| **Generation** | **0.224** | **0.031** | **7.33** | **<0.001\*** |
| Treatment (No Rescue) * Generation |  |  |  |  |
| **Treatment (Thermally Adapted Rescue) \***<br>**Generation** | **0.091** | **0.041** | **2.21** | **0.027\*** |
| Treatment (Non-adapted Rescue) * Generation | 0.055 | 0.041 | 1.33 | 0.185 |
| Random | 172 observations |  | Variance |  |
| ID:Thermal line | 58^ |  | 0.002 |  |
| Thermal line | 10 |  | 0.007 |  |

In generation one, no significant differences existed between any of the treatments (Non-Adapted - Adapted: Estimate = 0.015, SE = 0.057, *z* = 0.26, *P* = 0.794) (Table 2). In generations two and three, productivity in the rescue treatments differed significantly from the no rescue control, were not significantly different in generation 2 (Estimate = 0.095, SE = 0.057, *z* = 1.67, *P* = 0.095), but exhibited a significantly difference in generation 3 (Estimate = 0.097, SE = 0.045, *z* = 2.17, *P* = 0.030).

**Table 2:** Composite table of three GLMM results for each generation of the productivity of small, inbred, thermally adapted *T. castaneum* populations (Population size = 20, Experimental populations = 58 or 56) after receiving a rescue by a thermally adapted, or non-adapted, male rescuer or no rescue.

| | Gen 1<br>Estimate | SE | $z$ | $P$ | Gen 2<br>Estimate | SE | $z$ | $P$ | Gen 3<br>Estimate | SE | $z$ | $P$ |
| --- | --- | --- | --- | --- | --- | --- | --- | --- | --- | --- | --- | --- |
| Intercept<br>(No Rescue) | 5.810 | 0.055 | 105.84 | <2e-16 | 6.100 | 0.057 | 107.29 | <2e-16 | 6.263 | 0.051 | 122.93 | <2e-16 |
| Thermally<br>Adapted<br>Rescue | 0.061 | 0.058 | 1.05 | 0.291 | 0.284 | 0.058 | 4.90 | **<0.001\*** | 0.277 | 0.045 | 6.20 | **<0.001\*** |
| Non-Adapted<br>Rescue | 0.056 | 0.057 | 0.81 | 0.420 | 0.188 | 0.057 | 3.28 | **0.001\*** | 0.180 | 0.044 | 4.07 | **<0.001\*** |
|  | Gen 1<br>observation |  | Gen 1 variance |  | Gen 2<br>observation |  | Gen 2 variance |  | Gen 3<br>observation |  | Gen 3 variance |  |
| Random | 58 |  |  |  | 58 |  |  |  | 56 |  |  |  |
| ID:Thermal | 58 |  | <0.001 |  | 58 |  | 0.030 |  | 56 |  | 0.017 |  |
| Thermal | 10 |  | <0.001 |  | 10 |  | 0.005 |  | 10 |  | 0.016 |  |

## Discussion

We tested the disruptive effect of genetic rescue in locally adapted populations, by introducing thermally-adapted or outbred, non-adapted genetic rescuers to inbred thermally-adapted populations of *Tribolium castaneum* and then measuring productivity as a measure of population fitness. We show that populations receiving thermally-adapted rescue recovered fitness following two generations of inbreeding at a faster rate than populations that did not receive new genetic variation. When looking at each generation individually, both rescue treatments improved fitness in the second and third generations, when compared to the no-rescue control. In the third generation, fitness in populations that received thermally-adapted rescue improved over both the no-rescue control and non-adapted rescue treatments.

Introducing new genetic diversity into an inbred population is predicted to improve population fitness, and such genetic rescue effects have been observed in applied conservation studies (Madsen et al., 2020; Johnson et al., 2010). Here, we provide experimental support for this suggestion, as well as showing that the fitness of locally-adapted, inbred populations recovered at the greatest rate following the introduction of a rescuing individual from a population with a similar selective background. Importantly, we show that non-adapted rescuers also improved fitness compared with no rescue, despite the potential to disrupt adaptation, though the magnitude of the fitness increase was smaller and took longer to occur than when using a locally-adapted rescuer.

The ancestral population in our experiment is highly outbred and should represent an idea source of variation for use in genetic rescue (Ralls et al., 2020). Our findings show that local adaptations, evolved over 150 generations, considerably increased the efficacy of genetic rescue. Previous studies have suggested that the reinitiation of gene flow between populations following more that 20 generations of environmental divergence may risk outbreeding depression (Frankham et al., 2011). Our findings do not support this suggestion, adding to the growing body of evidence that the risk of outbreeding depression in genetic rescue may be exaggerated (Powell, 2023; Fitzpatrick and Reid, 2019; Fitzpatrick et al., 2015).

In our study, genetic rescue by an individual bearing local, thermal, adaptation was more effective than rescue by an outbred rescuer not bearing this adaptation. Our findings support recommendations that genetic rescue should utilise source populations inhabiting similar environments, reducing the risk of disrupting local adaptation (Lenormand, 2002). We used 10 independent thermally adapted lines in this study, all originating from the same genetically diverse ancestral population. As these populations adapted to 38°C independently (Vasudeva et al., 2019), they may represent differing subsets of genetic variation, providing alternative substrates for genetic rescue to act on. This design provides a proxy for population fragmentation, with subpopulations diverging from a larger outbred population. As the experimental evolution for thermal adaptation likely resulted in bottlenecking of the adapting populations (Dickinson, 2018), we predict these populations to be less genetically diverse than the outbred ancestral population.

In the thermally adapted populations we study, the key adaptation is the capacity to remain fertile while developing at 38°C (Sales et al., 2018; Vasudeva et al., 2019; Lewis, 2020). Our findings suggest that the introduction of outbred variation may have disrupted this adaptation. If the increase in fitness were solely attributable to the introduction of genetic variation, an outbred rescuer would generate the most impactive rescue effect (Ralls et al., 2020). We show that the addition of locally adapted variation was more important to rapid recovery from inbreeding effects, supporting the importance of rescue from populations with similar adaptive backgrounds (Edmands, 2007). Using translocations from locally-adapted populations could be important in a conservation context as they may improve population resilience to a changing climate (Fitzpatrick and Reid, 2019).

It is beyond the scope of this study to identify the causative genetic variation that conferred local adaptation, and that allowed locally adapted individuals to act as the most effective rescuers. Future work should aim to test if effective rescue is mediated by the transfer of beneficial alleles directly contributing to adapted traits (Kelly and Phillips, 2019), or by the purging of genetic load, mildly deleterious mutations, in genes associated with the adaptation (Grossen et al., 2020). Despite this lack of resolution, we use a powerful, highly replicated, experimental design to answer a question difficult to test in wild populations. Our findings are highly relevant to conservation scenarios and highlight the utility of experimental evolution in contributing to applied questions. We provide support for the consideration of local adaptation in genetic rescue programmes and suggest that well designed rescue regimes may be less prone to outbreeding depression than previously suggested.

## References

Al Hikmani, H., van Oosterhout, C., Birley, T., et al. (2024) Can genetic rescue help save Arabia’s last big cat? Evolutionary Applications, 17 (5). doi:10.1111/eva.13701.

Bachmann, J.C., Jansen Van Rensburg, A., Cortazar-Chinarro, M., et al. (2020) Gene Flow Limits Adaptation along Steep Environmental Gradients. The American Naturalist, 195. doi:10.5061/dryad.41ns1rn96.

Bartoń, K. (2024) MuMIn: Multi-Model Inference.

Bell, D., Robinson, Z., Funk, W.C., et al. (2019) The Exciting Potential and Remaining Uncertainties of Genetic Rescue. Trends in Ecology and Evolution, 34 (12): 1070–1079. doi:10.1016/j.tree.2019.06.006.

Bell, G. and Gonzalez, A. (2009) Evolutionary rescue can prevent extinction following environmental change. Ecology Letters, 12 (9): 942–948. doi:10.1111/j.1461-0248.2009.01350.x.

Bellard, C., Bertelsmeier, C., Leadley, P., et al. (2012) Impacts of climate change on the future of biodiversity. Ecology Letters. 15 (4) pp. 365–377. doi:10.1111/j.1461-0248.2011.01736.x.

Bijlsma, R., Westerhof, M.D.D., Roekx, L.P., et al. (2010) Dynamics of genetic rescue in inbred Drosophila melanogaster populations. Conservation Genetics, 11 (2): 449–462. doi:10.1007/s10592-010-0058-z.

Blomqvist, D., Pauliny, A., Larsson, M., et al. (2010) Trapped in the extinction vortex? Strong genetic effects in a declining vertebrate population. BMC Evol Biol, 10 (33). Available at: http://www.biomedcentral.com/1471-2148/10/33.

Brooks, M.E., Kristensen, K., Van Benthem, K.J., et al. (2017) glmmTMB Balances Speed and Flexibility Among Packages for Zero-inflated Generalized Linear Mixed Modeling.

Bryan, J. (2023) googlesheets4: Access Google Sheets using the Sheets API V4.

Charlesworth, D. and Willis, J.H. (2009) The genetics of inbreeding depression. Nature Reviews Genetics. 10 (11). doi:10.1038/nrg2664.

Clarke, J.G., Smith, A.C. and Cullingham, C.I. (2024) Genetic rescue often leads to higher fitness as a result of increased heterozygosity across animal taxa. Molecular Ecology. doi:10.1111/mec.17532.

Crnokrak, P. and Roff, D.A. (1998) Inbreeding depression in the wild. Heredity.

Dickinson, M. (2018) The impacts of heat-wave conditions on reproduction in a model insect, Tribolium castaneum. University of East Anglia.

Duval, C., Park, T., Miller, E.V., et al. (1939) Studies in Population Physiology. IX. The Effect of Imago Population Density on the Duration of the Larval and Pupal Stages of Tribolium. Ecology, 20 (3): 365–373.

Edmands, S. (2007) Between a rock and a hard place: Evaluating the relative risks of inbreeding and outbreeding for conservation and management. Molecular Ecology. 16 (3) pp. 463–475. doi:10.1111/j.1365-294X.2006.03148.x.

Fitzpatrick, S.W., Bradburd, G.S., Kremer, C.T., et al. (2019) Genetic rescue without genomic swamping in wild populations. *bioRxiv*. doi:10.1101/701706.

Fitzpatrick, S.W., Gerberich, J.C., Kronenberger, J.A., et al. (2015) Locally adapted traits maintained in the face of high gene flow. Ecology Letters, 18 (1): 37–47. doi:10.1111/ele.12388.

Fitzpatrick, S.W. and Reid, B.N. (2019) Does gene flow aggravate or alleviate maladaptation to environmental stress in small populations? Evolutionary Applications, 12 (7): 1402–1416. doi:10.1111/eva.12768.

Frankham, R. (2008) Genetic adaptation to captivity in species conservation programs. Molecular Ecology, 17 (1): 325–333. 10.1111/j.1365-294X.2007.03399.x.

Frankham, R. (2015) Genetic rescue of small inbred populations: meta-analysis reveals large and consistent benefits of gene flow. Molecular Ecology, 24 (11): 2610–2618. doi:10.1111/mec.13139.

Frankham, R. (2016) Genetic rescue benefits persist to at least the F3 generation, based on a meta-analysis. Biological Conservation, 195: 33–36. doi:10.1016/j.biocon.2015.12.038.

Frankham, R., Ballou, J.D., Eldridge, M.D.B., et al. (2011) Predicting the Probability of Outbreeding Depression. Conservation Biology, 25 (3): 465–475. doi:10.1111/j.1523-1739.2011.01662.x.

Grossen, C., Guillaume, F., Keller, L.F., et al. (2020) Purging of highly deleterious mutations through severe bottlenecks in Alpine ibex. Nature Communications, 11 (1). doi:10.1038/s41467-020-14803-1.

Haddad, N.M., Brudvig, L.A., Clobert, J., et al. (2015) Habitat fragmentation and its lasting impact on Earth’s ecosystems. Science Advances, 1 (2). doi:10.1126/sciadv.1500052.

Hartig, F. (2022) DHARMa: Residual Diagnostics for Hierarchical (Multi-Level / Mixed) Regression Models.

Hedrick, P.W., Adams, J.R. and Vucetich, J.A. (2011) Reevaluating and Broadening the Definition of Genetic Rescue. Conservation Biology. 25 (6) pp. 1069–1070. doi:10.1111/j.1523-1739.2011.01751.x.

Hedrick, P.W. and Garcia-Dorado, A. (2016) Understanding Inbreeding Depression, Purging, and Genetic Rescue. Trends in Ecology and Evolution. 31 (12) pp. 940–952. doi:10.1016/j.tree.2016.09.005.

Hedrick, P.W., Robinson, J.A., Peterson, R.O., et al. (2019) Genetics and extinction and the example of Isle Royale wolves. Animal Conservation, 22 (3): 302–309. doi:10.1111/acv.12479.

Hoegh-Guldberg, O., Jacob, D., Taylor, M., et al. (2018) 2018: Impacts of 1.5°C Global Warming on Natural and Human Systems. In: Global Warming of 1.5°C. An IPCC Special Report on the impacts of global warming of 1.5°C above pre-industrial levels and related global greenhouse gas emission pathways, in the context of strengthening the global response to the threat of climate change, sustainable development, and efforts to eradicate poverty.

Hope, R. (2022) Rmisc: Ryan Miscellaneous.

Howe, R.W. (1962) The effects of temperature and humidity on the oviposition rate of Tribolium castaneum (Hbst.) (Coleoptera, Tenebrionidae). Bulletin of Entomological Research, 53 (2): 301–310. doi:DOI: 10.1017/S0007485300048148.

Hufbauer, R.A., Szűcs, M., Kasyon, E., et al. (2015) Three types of rescue can avert extinction in a changing environment. Proceedings of the National Academy of Sciences, 112 (33): 10557. doi:10.1073/pnas.1504732112.

Ingvarsson, P.K. (2001) Restoration of genetic variation lost - the genetic rescue hypothesis. TRENDS in Ecology & Evolution, 16 (2).

Janus, M.C. (1989) Phenotypic diversity of Tribolium confusum pupae in heterogeneous environments. Entomologia Experimentalis et Applicata, 50 (3). doi:10.1111/j.1570-7458.1989.tb01203.x.

Johnson, W.E., Onorato, D.P., Roelke, M.E., et al. (2010) Genetic restoration of the Florida panther. Science, 329 (5999). doi:10.1126/science.1192891.

Kawecki, T.J. and Ebert, D. (2004) Conceptual issues in local adaptation. Ecology Letters. 7 (12) pp. 1225–1241. doi:10.1111/j.1461-0248.2004.00684.x.

Kelly, E. and Phillips, B.L. (2019) Targeted gene flow and rapid adaptation in an endangered marsupial. Conservation Biology, 33 (1): 112–121. doi:10.1111/cobi.13149.

King, C.E. and Dawson, P.S. (1972) “Population Biology and the Tribolium Model.” In Evolutionary Biology. doi:10.1007/978-1-4757-0256-9_5.

Laskowski, R., Radwan, J., Kuduk, K., et al. (2015) Population growth rate and genetic variability of small and large populations of Red flour beetle (Tribolium castaneum) following multigenerational exposure to copper. Ecotoxicology, 24 (5): 1162–1170. doi:10.1007/s10646-015-1463-3.

Lenormand, T. (2002) Gene flow and the limits to natural selection. Trends in Ecology & Evolution, 17 (4): 183–189. 10.1016/S0169-5347(02)02497-7.

Lewis, R.C. (2020) Thermal adaptation in a model pest insect. University of East Anglia. Available at: https://ueaeprints.uea.ac.uk/id/eprint/79830 (Accessed: 10 February 2025).

Loope, K.J., DeSha, J.N., Aresco, M.J., et al. (2024) Common-garden experiment reveals outbreeding depression and region-of-origin effects on reproductive success in a frequently translocated tortoise. Animal Conservation. doi:10.1111/acv.12977.

Lüdecke, D., Ben-Shachar, M., Patil, I., et al. (2021) performance: An R Package for Assessment, Comparison and Testing of Statistical Models. Journal of Open Source Software, 6 (60): 3139. doi:10.21105/joss.03139.

Macadam, A., Morgans, C., Cheok, J., et al. (2025) Assessing the potential for “assisted gene flow” to enhance heat tolerance of multiple coral genera over three key phenotypic traits. Biological Conservation, 306. doi:10.1016/j.biocon.2025.111155.

Madsen, T., Loman, J., Anderberg, L., et al. (2020) Genetic rescue restores long-term viability of an isolated population of adders (Vipera berus). Current Biology, 30 (21): R1297–R1299. 10.1016/j.cub.2020.08.059.

Onorato, D.P., Cunningham, M.W., Lotz, M., et al. (2024) Multi-generational benefits of genetic rescue. Scientific Reports, 14 (1). doi:10.1038/s41598-024-67033-6.

Palomares, F., Godoy, J.A., López-Bao, J.V., et al. (2012) Possible Extinction Vortex for a Population of Iberian Lynx on the Verge of Extirpation. Conservation Biology, 26 (4): 689– 697. doi:10.1111/j.1523-1739.2012.01870.x.

Pimm, S.L., Dollar, L. and Bass, O.L. (2006) The genetic rescue of the Florida panther. Animal Conservation, 9 (2): 115–122. doi:10.1111/j.1469-1795.2005.00010.x.

Pointer, M.D., Gage, M.J.G. and Spurgin, L.G. (2021) Tribolium beetles as a model system in evolution and ecology. Heredity. doi:10.1038/s41437-021-00420-1.

Posit team (2024) RStudio: Integrated Development for R.

Powell, D.M. (2023) Losing the forest for the tree? On the wisdom of subpopulation management. Zoo Biology. 42 (5) pp. 591–604. doi:10.1002/zoo.21776.

R Core Team (2024) R: A Language and Environment for Statistical Computing.

Ralls, K., Sunnucks, P., Lacy, R.C., et al. (2020) Genetic rescue: A critique of the evidence supports maximizing genetic diversity rather than minimizing the introduction of putatively harmful genetic variation. Biological Conservation. 251. doi:10.1016/j.biocon.2020.108784.

Rudin-Bitterli, T.S., Evans, J.P. and Mitchell, N.J. (2021) Fitness consequences of targeted gene flow to counter impacts of drying climates on terrestrial-breeding frogs. Communications Biology, 4 (1): 1195. doi:10.1038/s42003-021-02695-w.

Sales, K., Vasudeva, R., Dickinson, M.E., et al. (2018) Experimental heatwaves compromise sperm function and cause transgenerational damage in a model insect. Nature Communications, 9 (1). doi:10.1038/s41467-018-07273-z.

Sales, K., Vasudeva, R. and Gage, M.J.G. (2021) Fertility and mortality impacts of thermal stress from experimental heatwaves on different life stages and their recovery in a model insect. Royal Society Open Science, 8 (3). doi:10.1098/rsos.201717.

Skourti, A., Kavallieratos, N.G. and Papanikolaou, N.E. (2022) Demographic responses of Tribolium castaneum (Coleoptera: Tenebrionidae) to different temperatures in soft wheat flour. Journal of Thermal Biology, 103. doi:10.1016/j.jtherbio.2021.103162.

Sokal, R.R. and Sonleitner, F.J. (1968) The Ecology of Selection in Hybrid Populations of Tribolium castaneum. Ecological Monographs, 38 (4): 345–379.

Soule, M.E. (ed.) and Gilpin, M.E. (1986) Conservation biology: the science of scarcity and diversity. Sunderland, MA (USA) Sinauer Associates.

Tallmon, D.A., Luikart, G. and Waples, R.S. (2004) The alluring simplicity and complex reality of genetic rescue. Trends in Ecology and Evolution. 19 (9) pp. 489–496. doi:10.1016/j.tree.2004.07.003.

Turček, F.J. and Hickey, J.J. (1951) Effect of Introductions on Two Game Populations in Czechoslovakia. The Journal of Wildlife Management, 15 (1): 113–114. Available at: https://www.jstor.org/stable/3796784.

Vasudeva, R., Sutter, A., Sales, K., et al. (2019) Adaptive thermal plasticity enhances sperm and egg performance in a model insect. eLife. doi:10.7554/eLife.49452.001.

White, S.L., Rash, J.M. and Kazyak, D.C. (2023) Is now the time? Review of genetic rescue as a conservation tool for brook trout. Ecology and Evolution, 13 (5). doi:10.1002/ece3.10142.

Wickham, H. (2016) ggplot2: Elegant Graphics for Data Analysis.

Wickham, H., Averick, M., Bryan, J., et al. (2019) Welcome to the Tidyverse. Journal of Open Source Software, 4 (43): 1686. doi:10.21105/joss.01686.

Zajitschek, S.R., Zajitschek, F. and Brooks, R.C. (2009) Demographic costs of inbreeding revealed by sex-specific genetic rescue effects. BMC Evolutionary Biology, 9 (1). doi:10.1186/1471-2148-9-289.

